# Surviving heatwaves: thermal experience predicts life and death in a Southern Ocean diatom

**DOI:** 10.1101/2020.08.25.264028

**Authors:** Toby Samuels, Tatiana A. Rynearson, Sinéad Collins

## Abstract

Extreme environmental fluctuations such as marine heatwaves (MHWs) can have devastating effects on ecosystem health and functioning through rapid population declines and destabilisation of trophic interactions. However, recent studies have highlighted that population tolerance to MHWs is variable, with some populations even benefitting from MHWs. A number of factors can explain variation in responses between populations including their genetic variation, previous thermal experience and the intensity and duration of the heatwave itself. We disentangle the contributions of these factors on population survival and post-heatwave growth rates by experimentally simulating heatwaves (7.5 or 9.2 °C, for up to nine days) for three genotypes of the Southern Ocean diatom *Actinocyclus actinochilus*. The effects of simulated heatwaves on mortality and population growth varied with both genotype and thermal experience. Firstly, hotter and longer heatwaves increased mortality and decreased post-heatwave growth rates relative to milder, shorter heatwaves. Secondly, growth above the thermal optimum before heatwaves exacerbated heatwave-associated negative effects, leading to higher mortality during heatwaves and slower growth after heatwaves. Thirdly, hotter and longer heatwaves resulted in more pronounced changes to thermal optima (T_opt_) immediately following heatwaves. Finally, there is substantial intraspecific variation in mortality during heatwaves and in post-heatwave growth. Our findings shed light on the potential of Southern Ocean diatoms to tolerate MHWs, which will increase both in frequency and in intensity under future climate change.

## INTRODUCTION

Extreme temperature fluctuations in terrestrial and marine systems are occurring with higher frequency and duration than previously, and will increase further with continued anthropogenic climate change (Frölicher *et al*., 2018, Lyon *et al*., 2019, Oliver *et al*., 2019, Rohini *et al*., 2019). Marine heatwaves (MHWs) are one such example of temperature fluctuations, and are defined as “discrete prolonged anomalous warm water events” (Hobday *et al*., 2016) that can result in rapid population declines and reduced ecosystem functioning (Frölicher & Laufkötter, 2018, Oliver *et al*., 2019, Smale *et al*., 2019). Recent work has uncovered a broad range of organismal response to MHWs, from negative to positive (Bartosiewicz *et al*., 2019, Britton *et al*., 2020, Pansch *et al*., 2018, Saha *et al*., 2019, Stuhr *et al*., 2017). Furthermore, varied responses among species to MHWs can lead to significant food web alterations in marine habitats (Jones *et al*., 2018, Peña *et al*., 2019, Piatt *et al*., 2020, Ryan *et al*., 2017, von Biela *et al*., 2019), illustrating that the responses of individual species will influence the resilience of entire marine ecosystems under global change.

The ecological impact of marine heatwaves in the Southern Ocean has not previously received as much attention as MHWs in Arctic, temperature or tropical locations. However, between 2002 and 2018 nineteen heatwave events were detected across the Southern Ocean (Montie *et al*., 2020) and MHWs are predicted to increase in frequency in the Southern Ocean in coming decades (Frölicher *et al*., 2018). Indeed, heatwaves recorded across Antarctica in the summer of 2019-2020 are likely to have significant implications for the Antarctic ecosystem (Robinson *et al*., 2020). These recent events highlight the urgent need to understand how Southern Ocean organisms respond to MHWs. Diatoms dominate phytoplankton blooms in the Southern Ocean, are important primary producers that support the Southern Ocean ecosystem and are major exporters of silica and carbon from surface waters to marine sediments (Deppeler & Davidson, 2017). Constant elevated temperature can affect both population dynamics and the nutritional value of Southern Ocean diatoms (Boyd *et al*., 2016), indicating that diatom responses to future warming could have significant implications for trophic interactions and biogeochemical cycling. These findings are echoed by a large body of literature that concludes that environmental change is affecting marine phytoplankton, and will continue to do so in the future (Collins *et al*., 2020). Despite this, only a handful of studies have directly investigated the effect of MHWs on diatoms under controlled laboratory conditions (Bedolfe, 2015, Feijão *et al*., 2018, Remy *et al*., 2017).

From a physiological perspective, two key mechanisms affect the responses of organisms to thermal extremes, a) the cellular stress response (Schroda *et al*., 2015) and b) acclimation, which can result in “heat hardening” (Bowler, 2005). The cellular stress response, defined as the upregulation of stress response genes including those that express heat shock proteins, enhances tolerance to stressful conditions (Schroda *et al*., 2015). Acclimation, or gradual phenotypic plasticity (Kremer *et al*., 2018), describes the effect of altered gene expression and epigenetic modifications to adjust a phenotype (e.g. growth rate) in response to an environmental change (Angilletta, 2009, Kronholm & Ketola, 2018). In phytoplankton, acclimation can alter organismal fitness in the new environment over several asexual generations, and is reversible if the environmental cue stops (Anning *et al*., 2001, Brand *et al*., 1981, Kremer *et al*., 2018). Across a wide variety of marine taxa, numerous studies have demonstrated that previous acclimation to elevated temperatures can enhance tolerance (heat hardening) when exposed to thermal extremes (Hughes *et al*., 2019, Magozzi & Calosi, 2015, Pansch *et al*., 2018, Sasaki & Dam, 2019, Scharf *et al*., 2015, Stuhr *et al*., 2017).

Across both terrestrial and marine taxa, physiological responses to elevated temperature depend on the intensity and duration of thermal conditions within the context of the organism’s thermal niche. For example, environmental warming that increases temperature below thermal optima, the temperature at which growth rate is maximised, can be beneficial by enhancing metabolic activity (Angilletta, 2009). However, environmental warming that raises temperatures near or above thermal optima induce a number of physiological stress responses (Leung *et al*., 2017, Low *et al*., 2018, Madeira *et al*., 2013, Viant *et al*., 2003) that depend upon the duration of the thermal extreme, ranging from acute (hours to days) to chronic (days to weeks) (Huey & Bennett, 1990). Energy and resource investment into the expression of acclimation and stress response genes, such as those that produce heat shock proteins, incur fitness costs (Geider *et al*., 2009, Kingsolver & Woods, 2016, Krebs & Feder, 1997, Viant *et al*., 2003) and if these are high they can limit responses to future environmental change (Sokolova *et al*., 2012). In the context of marine heatwaves, the impact of elevated temperatures will be dependent upon the thermal niche of the organisms present, which is subject to both interspecific and intraspecific variation (Boyd *et al*., 2013). Furthermore, the state of cellular condition before heatwaves has the potential to affect population resilience to heatwaves when they do occur (Ainsworth *et al*., 2016, Saha *et al*., 2019, Short *et al*., 2015, Siegle *et al*., 2018, Stuhr *et al*., 2017).

Several studies in marine organisms show that resilience to MHWs depend upon several factors, including temperatures experienced prior to the thermal extreme (Siegle *et al*., 2018) and temperature variability (Lugo *et al*., 2020, Saha *et al*., 2019, Stuhr *et al*., 2017), and that resilience varies between species (Lugo *et al*., 2020, Magozzi & Calosi, 2015, Saha *et al*., 2019). Given these factors, it is not surprising that studies have a range of findings. Stuhr et al. (2017) demonstrate that episodic heatwaves enabled maintenance of growth and activity in corals where chronic exposure reduced them. In contrast, Lugo et al. (2020) found that low-temperature peaks in a fluctuating thermal regime did not provide relief after elevated temperature exposure in sea stars. Although these studies suggest that thermal experience interacts with genetic variation to constrain responses to marine heatwaves, this has not been explicitly addressed in marine phytoplankton, including diatoms.

To understand how intraspecific variation and thermal experience interact in determining survival and subsequent population growth, we investigated the growth response of the Antarctic diatom *Actinocyclus actinochilus* to heatwaves. We examined the impact of heatwaves on growth rates using three thermal variables: 1) normal temperature, 2) heatwave temperature and 3) heatwave duration. We used three genotypes of *A. actinochilus,* all isolated from the Ross Sea, to assess the potential for intraspecific variation in resilience to heatwaves. Experimental populations were grown either below or above their thermal optima for three weeks until they reached stationary phase, whereupon they were subjected to heatwaves of 7.5 or 9.2 °C for 0, 1, 3, 6, or 9 days, during which we measured mortality. Directly after heatwave exposure, we produced acute thermal performance curves for each genotype.

We examined whether previous acclimation at high temperatures dampens the negative consequences (increased mortality and/or decreased post-heatwave growth rates) of heatwave exposure by “heat hardening”, or if previous growth at a higher temperature exacerbates these negative effects which is consistent with deteriorating cellular condition rather than heat hardening. In addition to looking at how conditions prior to heatwaves affect survival and growth we show how heatwave survival and post-heatwave growth depend on heatwave intensity and length. We also investigate how heatwaves of different intensities and lengths alter the ability of populations to grow in different post-heatwave environments or respond to subsequent temperature changes by measuring acute plastic responses to temperature by characterising shifts in thermal optima (T_opt_) immediately following the heatwave. Finally, we explored the evidence for intraspecific variation in these heatwave responses.

## MATERIALS AND METHODS

### Genotype isolation and culture maintenance

Seawater samples were collected from surface waters (20 m depth) of the Ross Sea, Antarctica in January 2017. Individual cells and chains of the centric diatom *A. actinochilus* were isolated using a stereomicroscope (Olympus, Center Valley, USA) and a pipette, washed in sterile seawater and then incubated at 2 °C in 1:10 F/2 medium under continuous light at 80-100 μmol photons m^−2^ s^−1^ in 24 and 48-well microtiter plates. Successfully cultivated isolates were then grown at 3 °C under constant light intensity (~50 μmol photons m^−2^ s^−1^, measured using a 2-pi sensor) and an aliquot transferred to F/2 medium (Guillard, 1975) every 3-4 weeks. The three isolates used in this study (A4, B7, D8) were collected from two geographic locations 1) 74.64° S, 157° W (A4), 2) 73.91° S, 151.1° W (B7, D8).

Cultured isolates were identified to species using the 18S rDNA sequence. Genomic DNA was extracted from filtered biomass using the DNeasy 96 Plant Kit (Qiagen, Hilden, Germany) following the manufacturer’s protocol, with additional lysis of biomass at 65°C for 10-20 minutes to increase yield. The 18S was amplified using a 15 μL reaction mixture containing 1-2 ng DNA, 1X colorless GoTaq master mix (Promega, Madison, USA), and 0.5 mol L-1 each of the universal 18SA and 18SB primers (Medlin *et al*., 1988) in a thermal cycler (Eppendorf AG 22331 Mastercycler, Hamburg, Germany) at 94°C for 2 min, 40 cycles of 94°C for 30 s, 60°C for 60 s, and 72°C for 2 min followed by 10 min at 72°C. PCR amplicons were then purified by ethanol precipitation (Zeugin & Hartley, 1985) and quantified by a Nanodrop 1000 (Thermo Scientific, Waltham, USA). Amplicons were sequenced unidirectionally using the 18SB primer either on a 3500XL Genetic Analyzer (Applied Biosystems, Foster City, USA) at the University of Rhode Island Genomics and Sequencing Center, or on an ABI 3730XL (Applied Biosystems, Foster City, USA)) at Yale University’s Keck DNA Sequencing Facility. Sequences were analysed using Genomics Workbench software, V9.0.1 (Qiagen, Hilden, Germany) and BLAST (Altschul *et al*., 1990).

### Experimental design

The effect of thermal experience on the survival and post-heatwave growth rates of *A. actinochilus* was assessed in an experiment comprised of three phases (Figure 1). During the acclimation phase experimental populations in stationary phase were inoculated into 25 mL fresh medium at low density (1:50 dilution of dense culture) and incubated for 19 days at either 2.5 or 5.8 °C. This period provided sufficient time (5-6 generations at a growth rate of ~0.3 divisions per day) for experimental populations to attain their previous density and reach carrying capacity. One millilitre volumes of these experimental populations were then aliquoted into individual wells of 48-well microtitre plates, which were then incubated on a thermal gradient block during the heatwave phase at one of two temperatures, 7.5 or 9.2 °C. This phase lasted between zero (control, no heatwave) and nine days, with samples being collected for live-dead staining at 0, 1, 3, 6 and 9 days to determine mortality during heatwaves. Furthermore, experimental populations exposed to 0, 3 and 9 day heatwaves were transferred into nutrient-replete media and allowed to grow at a range of temperatures (−2.2, −0.4, +1.5, +2.7, +4.7, +6.2 and +7.6 °C) on a thermal gradient block to produce acute thermal performance curves in the final phase.

**Figure 1.**
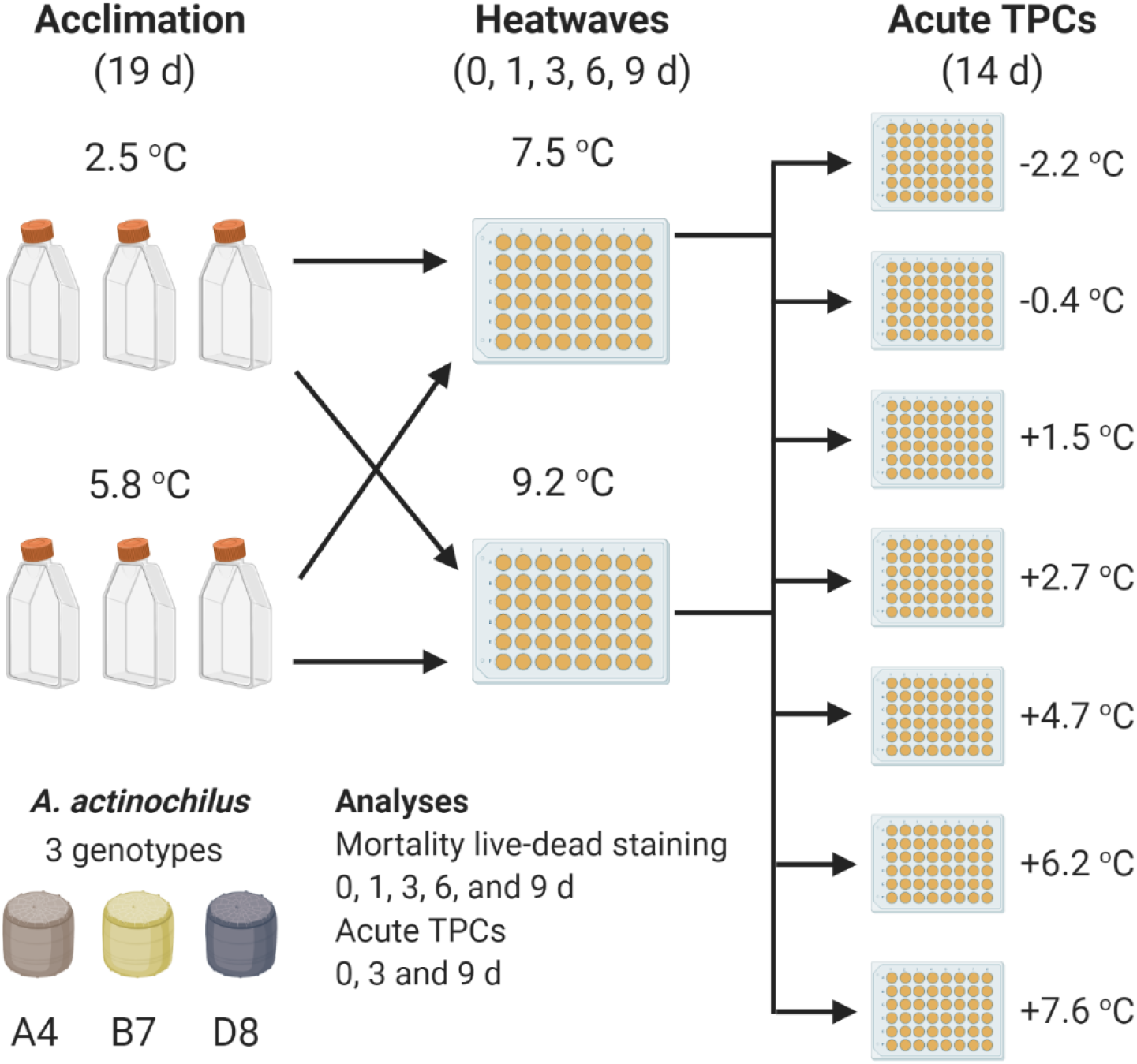
Overview of experimental design.

Heatwave temperatures (7.5 and 9.2 °C) were chosen to simulate thermally stressful environments for *A. actinochilus,* in which temperatures are either near to or beyond thermal maxima (Low *et al*., 2018). These temperatures, although not currently experienced in the Ross Sea where the genotypes used in this study were isolated, can occur in other parts of the Southern Ocean during heatwave events (Montie *et al*., 2020), and as such could be experienced by this species more broadly.

Experimental populations were diluted (1:20) into fresh media and allowed to grow until reaching stationary phase. Growth rates measured in this study are thus acute, rather than acclimated growth rates (Schulte *et al*., 2011). This enabled us to explore the role of recent thermal experience to extreme environmental fluctuations, when large temperature changes occur over days rather than weeks; our setup reflects immediate recovery from a heatwave rather than the acclimated growth usually used in laboratory experiments aimed at identifying the thermal niche of organisms (Boyd *et al*., 2013).

A full factorial design was implemented for three genotypes, two acclimation temperatures, and two heatwave temperatures. Mortality was measured for all heatwave durations (0,1,3,6 and 9 days). Post-heatwave growth rates were determined for heatwave durations of 0, 3 and 9 days only. All factorial combinations were grown as three independent triplicate experimental populations.

An inherent limitation of this experiment arises due to the known interaction between thermal stress and nutrient limitation (Rhee & Gotham, 1981, Thomas *et al*., 2017). When stationary phase experimental populations were transferred from the first to the second phase of the experiment they were not diluted into fresh media, and as such were exposed to elevated temperature under nutrient limitation. This was necessary for accurate measurements of mortality in experimental populations during heatwaves. Growth needed to be arrested, or mortality would be confounded by the growth of new cells.

We have accounted for this in two ways. Firstly, for each heatwave duration (0-9 days), each experimental population with differing thermal experiences was exposed to nutrient limitation for the same length of time. As such, differences in mortality and post-heatwave growth rates between treatment groups after a specific heatwave duration are directly comparable. Secondly, to quantify the effect of nutrient limitation on mortality in the experiment, we performed control treatments that maintained constant temperatures between the first and second phase of our experimental design, so that they were exposed to nutrient limitation only, without heatwaves. Experimental populations incubated at 2.5 or 5.8 °C in flasks in the first phase were transferred to roughly equivalent temperatures (2.7 and 6.2 °C) for the duration of the second phase, respectively. Due to space limitation on the thermal gradient blocks, it was not possible to perform both the heatwave treatments and control treatments simultaneously. As such, we treated these two data sets separately in our statistical analyses and compared the results of those analyses to infer the effect of heatwave temperatures. The survival analysis in this treatment was potentially confounded by the growth of new cells, so changes in density (both live and dead cells) over time were calculated to observe any potential growth.

### Incubation conditions

Experimental populations were incubated in a cooled incubator (Panasonic MIR-154) set to 3 or 6 °C for the first phase of both experiments. Temperature within flasks was measured in triplicate on four occasions, with the temperatures in the 3 and 6 °C incubators averaging 2.50 ± 0.24 °C and 5.77 ± 0.13 °C, respectively. Incubation during the heatwave and post-heatwave growth phases were performed on two thermal gradient blocks. For each block, two coolant circulators were used to cool/heat and then pump water/anti-freeze through either end of an aluminium block, on which culture vessels were incubated. The thermal gradient block was covered with sheets of insulation foam, with slots allowing microtitre plates to be in direct contact with the block surface. To enhance thermal conductivity between the block and the plates, custom cut aluminium sheets inserted into the base of microtitre plates, eliminating the air gap between the base of the plate and the bottom of the wells. Temperature values for each column were determined by taking measurements in 1 mL volumes of seawater within 48-well microtitre plates, with six wells recorded per measurement with a minimum of three measurements taken. Simulated heatwave temperatures (mean average +/− standard deviation) were 7.45 ± 0.34 and 9.24 ± 0.51. Temperatures included in analyses for the TPCs were −2.24 ± 0.66, −0.36 ± 0.40, +1.53 ± 0.43, 2.73 ± 0.46, 4.68 ± 0.28, 6.19 ± 0.4, 7.6 ± 0.42. Simulated heatwave temperatures for the control treatment were 2.73 and 6.19 °C. Lighting was provided from above using cool white aquarium LED lights, and light intensity was maintained at 45-55 μmol m^−2^ s^−1^, measured using a 2-pi sensor.

### Mortality analysis

On each of the heatwave analysis days (0, 1, 3, 6 and 9), samples of either 1 mL (from flask cultures) or 0.5 mL (from microtiter plates, diluted to a volume of 1 mL) were obtained from experimental populations after heatwave exposure, aliquoted into micro-centrifuge tubes and stained with Evans Blue dye at a final concentration of 0.02 %. Stained samples were incubated at 2.5 °C for a minimum of 1 hour. Samples were kept on ice until imaging (performed within 8 hours of staining). The order of sample imaging was randomised. This method was adapted from a previously published protocol (Garrison & Tang, 2014). Samples were loaded onto 1 mL Sedgewick rafter chambers and images acquired at 100 x magnification using a mounted EOS 800D Canon digital SLR camera. Images were taken to obtain counts per sample of >600 cells total, or until images of 300 squares (300 μL) were acquired. Live cells were determined as ranging in colour from dark to golden brown. Dead cells were determined as ranging in colour from light to dark blue.

### Determination of acute post-heatwave growth rates – Thermal performance curves (TPCs)

Acute growth rates of experimental populations, both before and after heatwave exposure, were determined at seven temperatures from −2.2 to +7.6 °C in roughly 1.5 °C intervals on a thermal gradient block. These temperatures broadly covered the focal organisms thermal tolerance range, in order to quantify effects on the growth response of experimental populations after heatwaves. After d0 (before heatwave), d3 and d9 of heatwave exposure, single technical replicates from each experimental population were inoculated from the acclimation (2.5 or 5.8 °C) or heatwave temperatures (7.5 or 9.2 °C) directly to temperatures ranging from −2.2 to 7.6 °C in 48-well microtitre plates and incubated for two weeks. Growth was monitored daily by measuring chlorophyll-a fluorescence on a Tecan Spark© plate reader, with experimental populations mixed using a pipette every second day to disperse cellular aggregates.

### Data and statistical analysis

All data analyses were performed in the R statistical environment (Team, 2019) and plots were produced in the package *ggplot2* (Wickham, 2016). Growth rates were calculated from chlorophyll-a fluorescence data using the equation below, where x_1_ and x_2_ are the estimated chlorophyll fluorescence values at the beginning (t_1_) and end (t_2_) of the fitted regression through the exponential phase of growth curves.

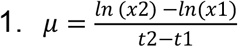

The maximum slope gradient was estimated from the growth curves using a sliding window approach across the two-week growth period, with the window providing the highest estimated value of growth rate accepted. The window length in all growth rate analyses was seven points, taken across consecutive days with the exception of anomalous data points removed from some growth series.

All estimated growth rates taken from growth curve fitted regressions were assessed using R^2^ confidence values. Fit confidence of growth rates varied both with temperature and growth rate (Supplementary figure 1), and varied most at the lowest TPC growth temperature (−2.24 °C) and at lower growth rate values (μ<0.2 d^−1^). Estimation of growth rates at extreme low temperatures and/or slower growth is more difficult due to small values of, and changes in, experimental population size. However, visual assessment of all growth rate fits was used to verify estimates and to confirm low, positive growth rates. For this reason, we chose not to exclude growth rates based upon an R^2^ value cut-off. Of the 598 positive growth rates, 589 had R^2^ values above 0.4, and 518 had R^2^ values above 0.75. The majority of growth rate estimates with R^2^ values below 0.75 were from the lowest (−2.24 °C, 34/80) and second lowest (−0.4 °C, 31/80), while eight out of nine growth rate estimates with R^2^ values below 0.4 were at −2.24 °C (Supplementary figure 1).

Survival data was analysed using generalized linear models with binomial distributions in the *glm* function, TPC growth rate data was analysed using general linear mixed models in the *lmer* function (package *lme4* (Bates *et al*., 2014)) and T_opt_ data was analysed within a general linear model using the *lm* function. In all models, heatwaves were defined by a term that multiplied their intensity (°C) with their duration (days), “cumulative heatwave intensity” (°C d), which was used as the sole variable for heatwave treatment due to collinearity between the two individual variables. For analysis of the survival data, genotype, acclimation temperature, cumulative heatwave intensity and percentage of live cells at day zero were used as predictor variables, with proportional relative survival (d_0_ = 1) as the response variable. For the analysis of growth data, genotype, growth temperature (both as a linear and a quadratic function), acclimation temperature and cumulative heatwave intensity were used as predictor variables, with growth rate as the response variable. TPC phase plate ID was included as random effect within this model, as ninety experimental populations were split across two plates per temperature. In the analysis of T_opt_, acclimation temperature, cumulative heatwave intensity and genotype were used as predictor variables, with T_opt_ values as the response variable. Post-hoc testing was performed using the emmeans package *emmeans()* function with Tukey tests to assess the significance of pairwise treatment comparison.

Relative growth differences (g) of experimental populations exposed to heatwaves after three or nine days (d_x_) of heatwave exposure, relative to the growth of experimental populations not exposed to heatwaves (d_0_), was calculated using the following equation:

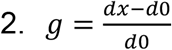

In order to assess shifts in thermal optima (T_opt_), the temperature at which growth rate is maximised. Although a number of thermal traits can be derived from fitted TPCs, T_opt_ has been shown in numerous studies to be highly responsive to the effect of recent thermal experience (Bernhardt *et al*., 2018, Kremer *et al*., 2018, Padfield *et al*., 2016, Staehr & Birkeland, 2006). All thermal performance curves were fit using a modified Norberg function (Thomas *et al*., 2012) within the growthTools package (Kremer, 2020), using the function *get.nbcurve.tpc()*. Curves were fit from growth data across the thermal gradient for each biological replicate, allowing triplicate curves to be produced per treatment, creating ninety curves in total. Fits were assessed for goodness-of-fit visually, with poor fits removed from the analysis. Eight fits were removed due to this assessment, with the remaining fits having R^2^ values > 0.4 (78 fits had R^2^ values >0.8). All data for genotype D8 acclimated to 5.8 °C and then exposed to 9.2 °C heatwaves for nine days were removed, due to erratic growth. Furthermore, due to enhanced growth at the lower thermal extremes in the experimental populations of genotype A4 acclimated at 2.5 °C and then exposed to 7.5 °C heatwaves for three days, the relationship between growth and temperature was no longer quadratic, and as such all data for this treatment was also removed from the analysis.

## RESULTS

### Population survival during heatwaves

The interaction of acclimation temperature with heatwave temperature and duration on the survival of experimental populations varied between genotypes (A4, B7 and D8), but general trends could be identified (Figure 2). Acclimation at either 2.5 or 5.8 °C for 19 days had consequences for cell survival before heatwave exposure. The average percentage of live cells at day zero for each genotype was 95-98 % for experimental populations acclimated at 2.5 °C, and 86-93 % for experimental populations acclimated at 5.8 °C. Experimental populations of genotype B7 acclimated to 2.5 °C had the highest percentage of live cells before heatwaves (98 %), while experimental populations of genotype D8 acclimated to 5.8 °C had the lowest (86 %). In a general linear model with the percentage of live cells at day zero as a response variable, acclimation temperature (F=45.03, df=1, p<0.001), genotype (F=4.47, df=2, p<0.035) and the interaction between them (F=4.55, df=2, p<0.034) were all significant.

**Figure 2.**
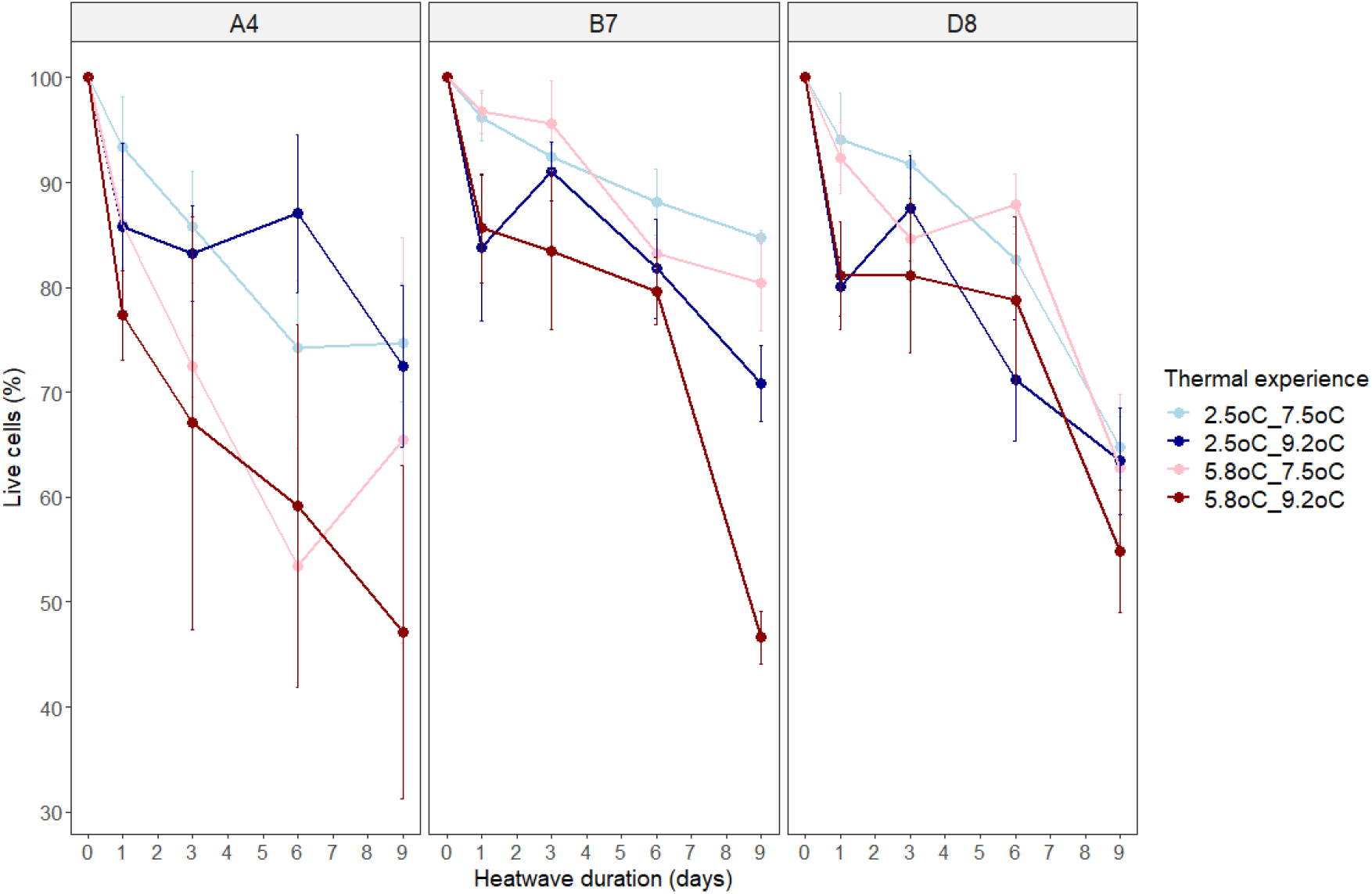
Survival of experimental populations during simulated heatwaves, shown as the percentage of live cells in experimental populations relative to the percentage of live cells on day 0. Thermal experience is defined by the combination of acclimation temperature (2.5 or 5.8 °C) and heatwave temperature (7.5 or 9.2 °C). Panels show the results for each genotype (A4, B7 and D8) of *Actinocyclus actinochilus*. Data points represent average values for three independent replicates ± one standard deviation.

Survival declined with increasing heatwave duration in all experimental populations, and the size of this effect varied both by genotype and acclimation temperature (figure 2). By day nine, the average percentage of live cells remaining in the population had declined to between 47 and 85 % relative to day zero values. For two of three genotypes, A4 and B7, experimental populations grown at 2.5 °C in the first phase of the experiment survived better when exposed to heatwaves compared to experimental populations grown at 5.8 °C. In the third genotype, D8, survival declined consistently regardless of acclimation temperature (figure 2).

We found evidence consistent with warmer heatwaves (9.2 °C, darker lines) decreasing survival more than cooler heatwaves (7.5 °C, lighter lines) for genotypes B7 and D8 (figure 2). In a binomial generalised linear model (GiLM), the effects of cumulative heatwave intensity (°C d) (GiLM, χ^2^=12.3685, df=1, p=0.0004) and acclimation temperature (GiLM, χ^2^=3.4029, df=1, p=0.0651) explain survivorship in these treatments. Cumulative heatwave intensity reduced survival in all three genotypes, with values declining to 40-60 % after nine days of the warmer heatwave. Experimental populations acclimated to 2.5 °C had lower mortality during heatwaves than those acclimated to 5.8 °C (Figure 2). There was no statistical evidence for genotype influencing survival (GiLM, χ^2^=1.8158, df=2, p=0.4034). Addition of interactions between response variables reduced model AICc values, and none of these terms were significant (Supplementary table 1).

In the control treatment, experimental populations were maintained in a stable thermal environment for up to 9 days to determine if time spent in stationary phase alone impacted survival, even without heatwaves. Of the six control treatment groups (genotypes A4, B7 and D8, in two thermal regimes), we only able to collect data for five, due to culturing failure in experimental populations of genotype D8 acclimated at 5.8 °C. In general, experimental populations in the control treatment maintained high percentages of live cells relative to the percentage of live cells on day zero (Supplementary figure 3). After nine days, percentages of live cells relative to the percentage of live cells on day zero ranged between 88-99 % in four of the five successfully cultivated control treatment groups. This in contrast to experimental populations exposed to nine days of heatwave treatment, that had percentages of live cells relative to the percentage of live cells on day zero ranging between 47-85 % (Supplementary figure 3). It should be noted that in the fifth control treatment group, genotype D8 acclimated at 5.8 oC and then exposed to 6.2 oC for nine days, survival declined more substantially (70 % after nine days). However, in a statistical analysis of all five control treatment groups, the effect of cumulative heatwave intensity (°C d^−1^) was non-significant (GiLM, χ^2^=0.3944, df=1, p=0.53) (Supplementary table 2), in contrast to the significance of this term in the heatwave treatment (GiLM, χ^2^=12.3685, df=1, p=0.0004) (Supplementary table 1). These differences indicate that the heatwave intensities used in the main experiment (7.5 and 9.2 °C) caused significant declines in the percentage of live cells that in general were not experienced at control treatment temperatures (2.7 and 6.2 °C) (Supplementary figure 3).

Density (cells/mL, total live and dead cells) of sampled experimental populations for both the heatwave and control treatments remained relatively constant (Supplementary figures 2 and 4). Minor fluctuations in density occurred, but these were not the large increases that would be associated with sustained growth. This indicates that the observed decline in live cells within experimental populations was not confounded by growth in both experiments.

### Thermal performance curves before heatwaves

We characterised the plastic responses of experimental populations to changes in temperature before heatwaves with thermal performance curves, using assay temperatures between −2.2 to +7.6 °C. There was intraspecific variation both in plasticity, defined as change in growth rate with increasing temperature below T_opt_, and in the effect of previous growth temperature (acclimation) on growth rates (Figure 3, left-hand panels). For example, genotype A4 showed limited plasticity from the lowest temperature assayed (−2.2 °C, μ = 0.37 +/− 0.02) to the temperature at which the highest growth rate was recorded (+4.7 °C, μ = 0.49 +/− 0.02), when previously grown at 3 °C (top-left panel, blue line). In contrast, genotype B7 was more plastic, with growth rate increasing by more than three-fold from −2.2 °C (0.11 +/− 0.01) to 2.7 °C (0.35 +/− 0.02) (middle-left panel, blue line). For genotypes A4 and B7, the effect of previous growth temperature had minimal influence on plasticity (blue and red lines substantially overlap in top-left and middle left panels). However, previous growth at 5.8 °C for genotype D8 resulted in significantly decreased growth rates across the central and upper temperature range, compared to experimental populations grown previously at 2.5 °C (bottom-left panel).

**Figure 3.**
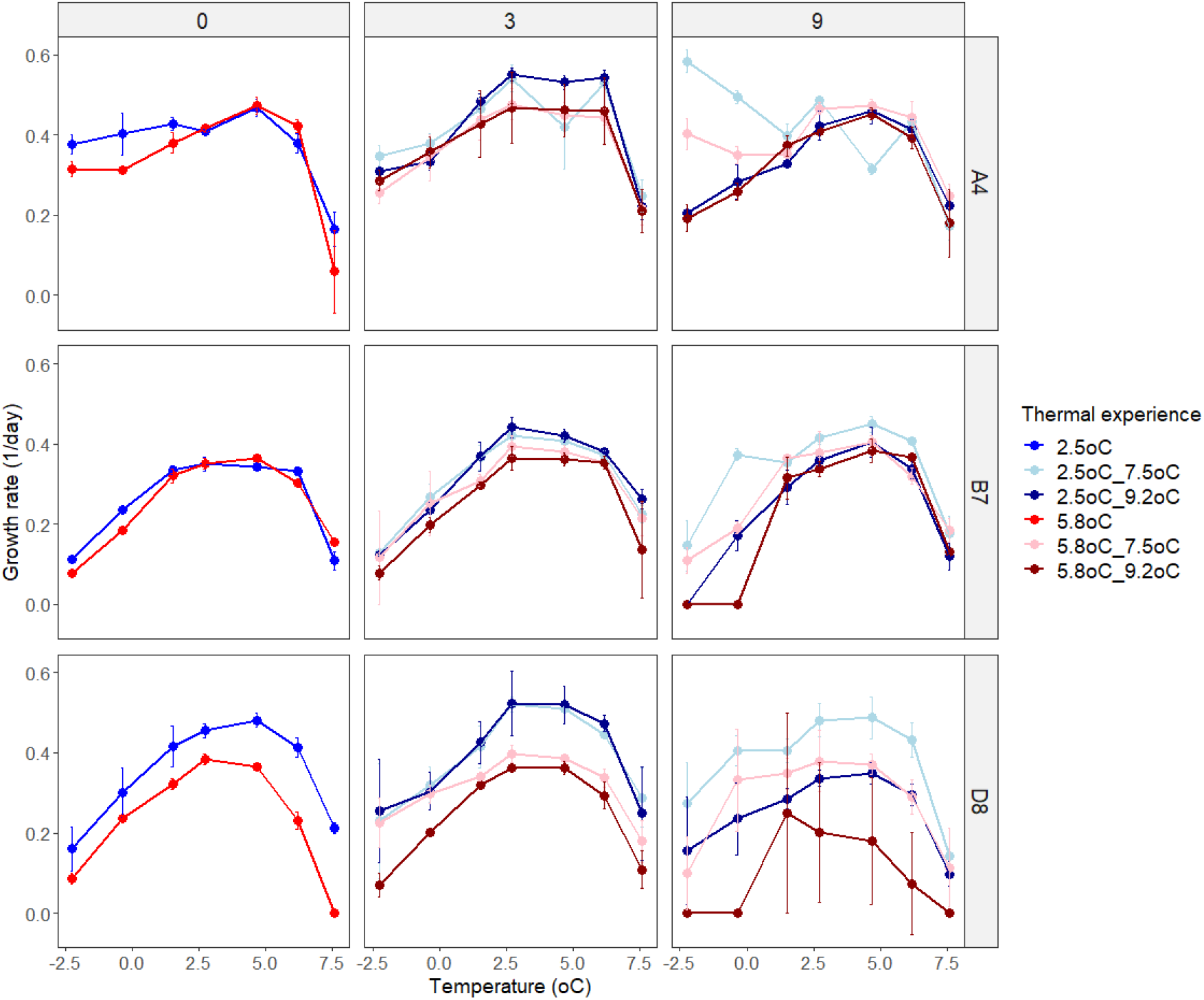
Maximum growth rate of experimental populations at seven temperatures before (day 0) and after heatwaves of 3 or 9 days in duration, for each genotype. Thermal experience combines acclimation temperature (2.5 or 5.8 °C) and heatwave temperature (7.5 or 9.2 °C; no temperature noted for cases with no heatwave). Data shown are average values across three replicates ± one standard deviation.

### Growth post-heatwave exposure

To assess the effect of heatwaves on plastic responses to warming, we constructed a second set of thermal performance curves for each experimental population following the heatwaves (Figure 3). Exposure to heatwaves of four differing cumulative intensities (7.5 or 9.2 °C, for three or nine days) resulted in both increases and decreases in growth rates relative to experimental populations not exposed to heatwaves (Figure 4). We explored the factors affecting changes in growth using a general linear mixed model (GLMM, supplementary table 3).

**Figure 4.**
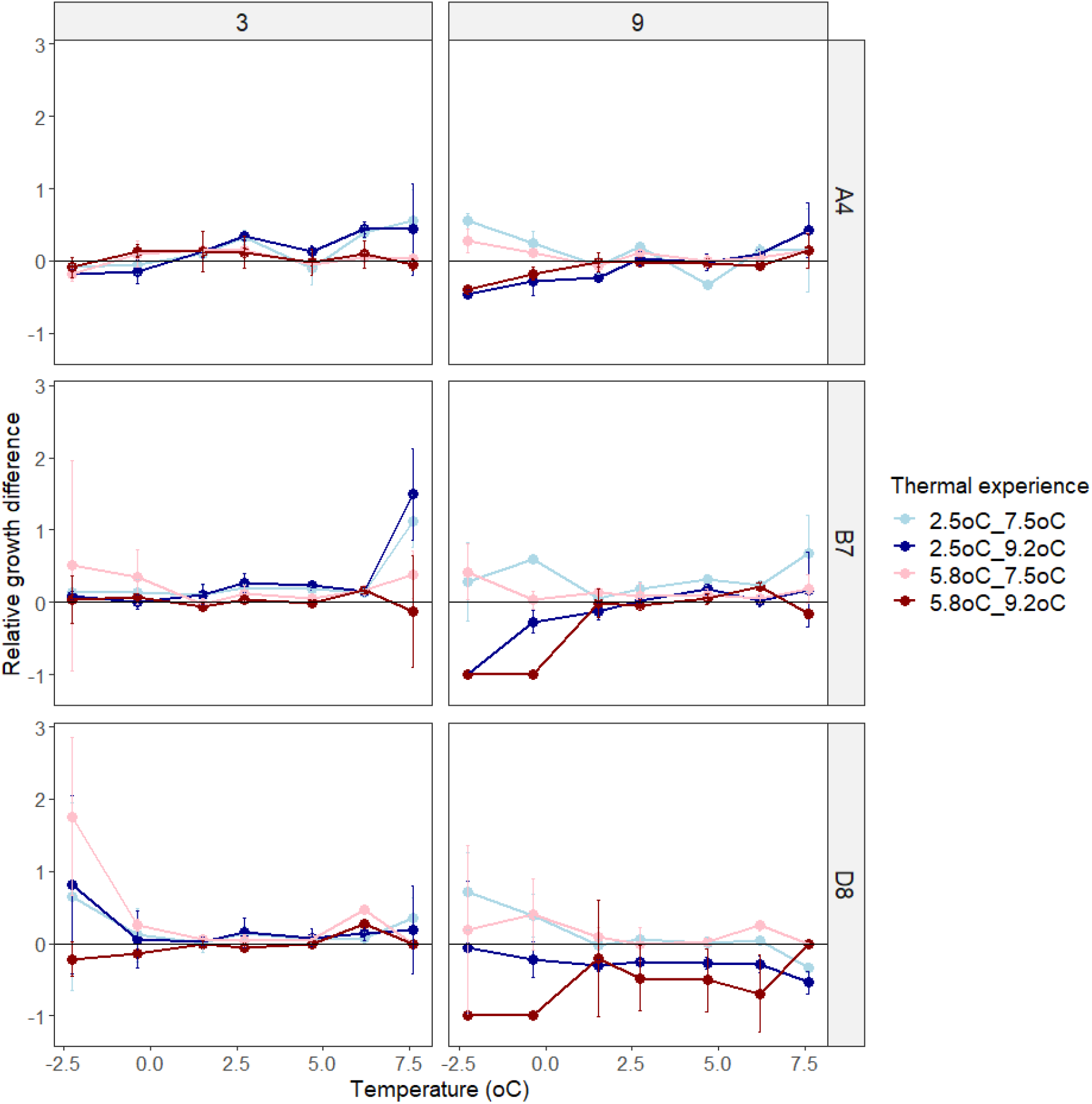
Relative growth difference of experimental populations grown at seven temperatures after exposure to 3 or 9 day heatwaves compared to experimental populations not exposed to heatwaves, for each genotype. Data represent average values across three replicates ± one standard deviation.

Since not all temperature response curves had the same shape, we described the relationship between temperature and growth rate using a linear function (GLMM, F=71.7293, df=1, p<0.001) and a quadratic function (GLMM, F=76.0014, df=1, p<0.0001). Together, these terms capture the shape of the temperature response curves and explain 21 % of the total variation explained by the model. Genotype (GLMM, F=239.8280, df=2, p<0.0001) and acclimation temperature (GLMM, F=133.9460, df=1, p<0.0001) together explained the majority (53 %) of the variance in growth rate (Figure 3, supplementary table 3). Here, experimental populations that had been acclimated at a lower temperature generally grew faster at a given temperature than those acclimated at a higher temperature (Figure 3, central and right hand panels).

The combined influence of heatwave temperature and duration was analysed using cumulative heatwave intensity (°C d, see methods), which had a large and significant effect on experimental population growth (GLMM, F=67.8707, df=4, p<0.0001). Post-hoc testing (Supplementary tables 4a-c) revealed that the effect of cumulative heatwave intensity was not linear. In general, growth was unaffected or enhanced by mild heatwaves, but reduced by intense heatwaves, with the extent of this modulated by genotype and acclimation temperature. For example, the most extreme growth rate increases occurred in B7 acclimated to 2.5 °C, where all but the most intense heatwaves (79.8 °C d, 9.2 °C for nine days) significantly increased growth rates relative to pre-heatwave values (Figure 4, Supplementary table 4b). However, the most intense heatwaves (79.83 °C d, 9.2 °C for nine days) significantly decreased growth rate, relative to both pre-heatwave values and to experimental populations exposed to milder heatwaves, in almost all pairwise comparisons across genotypes (Supplementary tables 4a-c). After intense heatwaves, growth at lower temperatures was either significantly reduced or completely arrested in all three genotypes. Furthermore, genotype D8 exposed to 9.2 °C heatwaves decreased across the entire range of temperatures measured, indicating that elevated temperatures globally reduced growth rates, even when experimental populations were returned to ambient temperatures.

A number of interactions between all individual variables within the main growth rate statistical model were also identified (supplementary table 3), but these had smaller effect sizes than the individual variables themselves, so only three interactions will be examined in detail here. First, the interaction with the largest effect size was genotype with growth temperature (GLMM, F=56.2742, df=2, p<0.0001), which can be explained by the differences in plasticity between genotypes described above.

Second, growth temperature also interacted with cumulative heatwave intensity (GLMM, F=17.7256, df=4, p<0.0001). Growth at both temperature range extremes were the most affected by cumulative heatwave intensity, with less intense heatwaves elevating growth rates while more intense heatwaves reduced them. Very high cumulative heatwave intensities (nine-day heatwaves of 9.2 °C) resulted in the largest reductions in growth rate, particularly at low temperatures (Figure 4, right-hand panels). For genotypes B7 and D8, growth at both −2.2 and −0.4 °C was completely inhibited after a thermal experience where experimental populations had been grown at 5.8 °C, and then exposed to a 9.2 °C heatwave for nine days. Finally, acclimation temperature also interacted with genotype (GLMM, F=12.8787, df=2, p<0.0001).

Due to the lower confidence of growth rate fits at the extreme low temperature (−2.24 °C), the statistical analysis of growth rates was repeated but without data from this temperature (Supplementary table 5). Despite some minor changes in relative effect sizes and the significance of low-effect interactions, the relative size and significance of major terms and interactions remained constant. As such, we decided to continue to use the full dataset in further analyses of the experiment.

Differences in thermal optima (T_opt_) between experimental populations were observed and were dependent upon thermal experience (Figure 5). Both acclimation temperature and heatwave temperature resulted in shifts of T_opt_ in all three genotypes for at least some of the heatwave durations. In general, hotter heatwaves resulted in a higher T_opt_. Over time, the difference in T_opt_ between hotter and cooler heatwaves either held steady or grew, but did not get smaller. The magnitude of the effects of acclimation temperature, heatwave temperature, and heatwave duration on T_opt_ were genotype specific. For example, before heatwave exposure, experimental populations of A4 acclimated at 5.8 °C had a T_opt_ higher than those acclimated at 2.5 °C. In comparison, genotype B7 had the same T_opt_ regardless of acclimation temperature (Figure 5). Heatwaves raised T_opt_ values in genotype A4, regardless of acclimation temperature, but the duration and temperature of a heatwave required to do this depended on acclimation temperature: both 7.5 and 9.2 °C heatwaves caused an upward shift in T_opt_ for experimental populations acclimated to 2.5 °C (Figure 5, top left), but only 9.2 °C for 9 days was able to further increase T_opt_ values for those acclimated at 5.8 °C (Figure 5, top right). In contrast, heatwaves caused no significant shifts in T_opt_ in genotype D8, regardless of heatwave intensity or duration (Figure 5, bottom panels).

**Figure 5.**
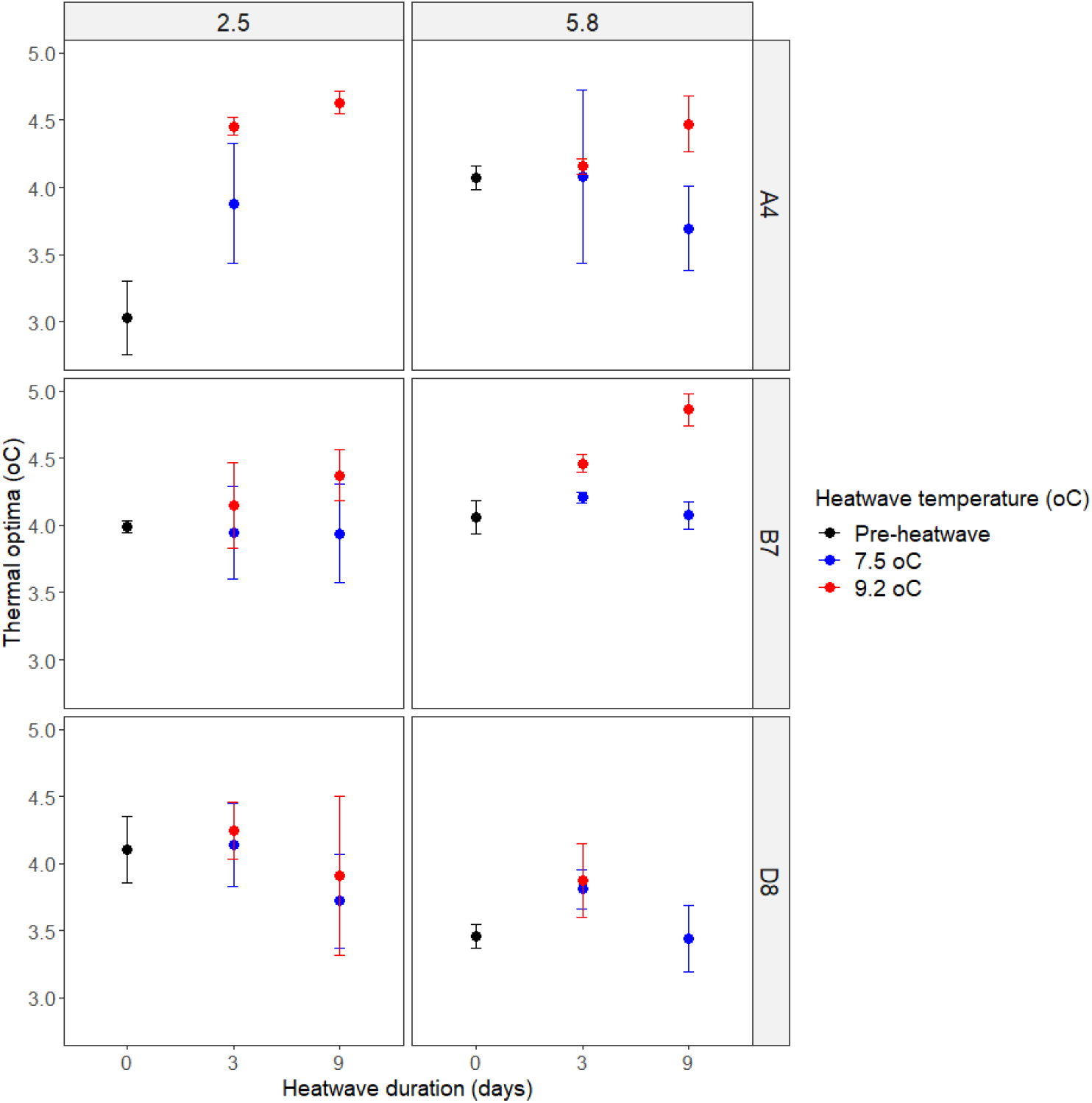
Growth rate thermal optima (°C) of experimental populations before (black) and after (red/blue) heatwave exposure. Thermal performance curves were produced before heatwaves (d0) and after 3 and 9 days of heatwave exposure, for different acclimation temperatures (columns) and genotypes (rows). Error bars represent ± one standard deviation across three biological replicates.

In a statistical model, cumulative heatwave intensity (GLM, df=4, F=12.08, p<0.0001) and genotype (GLM, df=2, F=8.08, p<0.0001) significantly explained variation in T_opt_, and significant interactions of genotype with both acclimation temperature (GLM, df=2, F=5.99, p=0.0041) and cumulative heatwave intensity (GLM, df=2, F=2.19, p=0.040) were identified (Figure 5, supplementary table 6). Post-hoc testing of the effect of cumulative heatwave intensity, when all other variables are kept equal, showed that heatwaves of 9.2 °C significantly increased T_opt_ relative to pre-heatwave experimental populations by 0.4 °C for three day (26.6 °C d, p<0.0001) and 0.5 °C for nine days (79.8 °C d, p<0.0001). In contrast, heatwaves of 7.5 °C did not significantly alter T_opt_ from pre-heatwave values, for either three day (21.9 °C d, p=0.2024) or nine day (65.6 °C d, p=0.9812) heatwaves.

## DISCUSSION

Previous studies have shown both negative and positive effects of heatwaves on marine organisms (Bartosiewicz *et al*., 2019, Lugo *et al*., 2020, Pansch *et al*., 2018, Roberts *et al*., 2019, Siegle *et al*., 2018, Stuhr *et al*., 2017). Here, we systematically disentangled how thermal experience and genotype affect the mortality and growth responses of the Southern Ocean diatom *Actinocyclus actinochilus* to simulated heatwaves, to show how both negative and positive heatwave effects can occur. We find that even under controlled laboratory conditions, a wide range of growth and mortality effects occur during and after exposure to heatwaves that can be explained by differences in temperature regimes: as expected, hotter and longer heatwaves cause higher mortality and affect post-heatwave growth more than more moderate heatwaves, and the extent of this depends on previous growth temperature. We also find that genotypes vary in their responses to heatwaves. Together, our data show that understanding the effects of marine heatwaves on primary producers requires taking both environmental and genetic context into account, and that while there are general patterns, a range of responses to heatwaves should be expected between populations with differing thermal histories.

More intense and longer heatwaves consistently resulted in higher mortality across all three genotypes (Figure 2, supplementary table 1). This supports previous work showing that cumulative thermal stress is a strong predictor of population decline in marine populations and ecosystems (Eakin *et al*., 2010, Marbà & Duarte, 2010, Pansch *et al*., 2018, Siegle *et al*., 2018, Stuhr *et al*., 2017). Accumulated stress beyond a critical threshold results in mortality due to the degradation of cellular components and disruption of physiological functioning (Feijão *et al*., 2018, Lesser, 2006, Magozzi & Calosi, 2015, Schroda *et al*., 2015). Although exposure to non-lethal temperatures can also increase tolerance to further stress in some marine organisms (Clapp *et al*., 1997, Magozzi & Calosi, 2015, Sasaki & Dam, 2019), we do not find evidence for this in our experiment. Instead, we find that experimental populations grown at 5.8 °C (above T_opt_) prior to heatwave exposure had higher levels of mortality than those grown at 2.5 °C (below T_opt_), in both the heatwave and control treatments. These results show that previous thermal experience exacerbated the negative consequences of heatwaves in this experiment. A number of studies have made similar conclusions; that previous acclimation to sub-lethal temperatures weakened the capacity to respond to further stress, rather than provide a “heat-hardening” effect (Marbà & Duarte, 2010, Pansch *et al*., 2018, Siegle *et al*., 2018). For example, Siegle et al. (2018) found that copepods isolated from splash pools with differing thermal histories responded differently to simulated heatwaves. Individuals that had experienced sub-lethal temperatures were less likely to survive simulated heatwaves, with this effect exacerbated by increasing heatwave intensity (Siegle *et al*., 2018). We suggest that in our study, the effect of previous heat exposure is related to whether experimental populations were acclimated above T_opt_, and nearer to the upper limit of temperatures normally experienced in the Southern Ocean during diatom blooms (5.8 °C), or below T_opt_, and within the normal temperature range for diatoms growing in the Southern Ocean (2.5 °C) (Boyd, 2019). Above T_opt_, one could reasonably suppose that cells would be stressed, and could accumulate damage, even if they are able to grow quickly in the short term, which is consistent with, for example, reactive oxygen associated with rapid growth due to CO2 enrichment (Lindberg & Collins, 2020). Below T_opt_, cells would be operating normally, and even if they are not under ideal conditions, it is unlikely that they are experiencing severe thermal stress.

We investigated the effect of thermal experience on growth immediately following heatwaves by comparing thermal performance curves before and after heatwave exposure. Previous studies on Southern Ocean diatoms (Andrew *et al*., 2019, Boyd *et al*., 2016, Boyd, 2019, Boyd *et al*., 2013, Coello-Camba & Agustí, 2017, Xu *et al*., 2014) largely present growth data for acclimated TPCs, where growth rate at a given temperature is recorded after several generations of acclimation to a stable thermal environment (Brand *et al*., 1981). In contrast, we have used acute thermal responses to investigate population dynamics that are rapidly altered by environmental fluctuations such as MHWs. Our rationale for investigating acute responses is that population dynamics, and the ecological consequences of them, immediately following heatwaves will be driven by acute responses to the post-heatwave temperatures, as ecological and evolutionary processes will not be paused while populations acclimate for several generations. We found that plastic responses to warming varied between genotypes and thermal experience. Exposure to heatwaves affected growth in all three genotypes (Supplementary table 3), but the extent of this impact substantially differed between them (Figure 3). Broadly, genotype A4 displayed the largest tolerance (low mortality, high growth rates) to simulated heatwaves, while genotype D8 displayed the lowest tolerance. Interestingly, three days of heatwave exposure did not have any significant negative consequences for growth in any genotype (Figure 4). This high tolerance to short but intense heatwaves was surprising, as a previous study has highlighted the negative impact of short heatwaves on diatoms; exposure to simulated heatwaves for three days significantly reduced growth rates in the model diatom *Phaeodactylum tricornutum* (Feijão *et al*., 2018). In contrast, we identified that three-day heatwaves sometimes had positive effects on growth rate at either thermal extreme in all three genotypes tested (Figures 2 and 3). An increase in growth rate at upper thermal extremes after heatwave exposure could indicate acclimation (Magozzi & Calosi, 2015, Scharf *et al*., 2016, Stuhr *et al*., 2017), in which growth at elevated temperatures cause rapid physiological alterations that limit the negative impact of stress at temperatures above T_opt_ (Low *et al*., 2018). However, we also saw high variation in growth within genotypes at the upper temperature range, indicating that acclimation at very high temperatures was variable (Figure 4). This may be because cells at very high temperatures are more stressed, and cell function is deteriorating idiosyncratically (Feijão *et al*., 2018, Magozzi & Calosi, 2015, Sørensen *et al*., 2013).

The most intense heatwaves (79.83 °C d, 9.2 °C for nine days) resulted in the largest reductions in growth rate observed in this study, in some cases completely arresting growth at thermal extremes (Figures 3 and 4). This is consistent with our mortality measurements, and other studies showing that as experimental populations reach a critical level of accumulated thermal stress growth rates and survival substantially declines (Marbà & Duarte, 2010, Pansch *et al*., 2018, Siegle *et al*., 2018). The largest reductions in survival occurred between six and nine days of heatwave exposure (Figure 2), indicating that differences in growth rates seen between three and nine day heatwave exposed experimental populations is likely driven by increased cellular stress (Figure 3). Strikingly, even after nine days of heatwave exposure, a subset of experimental populations displayed remarkable tolerance to elevated temperature, measured as growth rate (Figures 3 and 4). In genotypes A4 and B7, growth at 7.6 °C was enhanced after heatwave exposure, further supporting that acclimation can partially mitigate the negative effects of growth under elevated temperatures in some cases (Magozzi & Calosi, 2015, Scharf *et al*., 2016, Stuhr *et al*., 2017). Interestingly, some experimental populations exposed to nine day heatwaves also displayed enhanced growth rates at low temperatures (Figure 4). As with experimental populations exposed to shorter heatwaves, there was high variation within genotypes in lower temperature growth rates, which suggests that the effects of stress may be idiosyncratic in extreme cases, where the breakdown versus improvement of cellular function may have a high stochastic component.

Despite variation noted at low temperatures, there was still consistent enhanced growth at the extremes of the thermal range after heatwave exposure. We therefore hypothesize that thermal stress experienced at heatwave temperatures acted to enhance growth both at low (−2.2, −0.4 °C) and high (7.6 °C) thermal extremes (Figure 3). This idea is in line with previous studies that have used genomic and transcriptomic approaches to characterise the thermal adaptations of diatoms to the Southern Ocean, demonstrating that common stress mechanisms can be employed at both thermal extremes (Mock *et al*., 2017, Pargana *et al*., 2020). Growth of *Fragilariopsis cylindrus* under both thermal extremes (−2, +11 °C) revealed similarities in altered gene expression relative to ambient conditions, indicating that generic stress responses were potentially employed in both thermal environments (Mock *et al*., 2017), and *Leptocylindrus aporus* cultured at high and low temperatures revealed that the expression of heat shock proteins were upregulated under both thermal regimes (Pargana *et al*., 2020). The production of HSPs is known to increase during growth above T_opt_ (Leung *et al*., 2017, Rousch *et al*., 2004), and has been shown to be maximised as temperatures approach thermal maxima (Tmax) (Low *et al*., 2018). However, HSPs are also produced during cold shock responses, which is consistent with the observation that heat shocks can enhance tolerance to subsequent cold shocks (Burton *et al*., 1988, Goto & Kimura, 1998, Scharf *et al*., 2015, Scharf *et al*., 2016). Furthermore, metabolic activity scales with increasing temperature, resulting in elevated metabolism in experimental populations exposed to higher temperatures (Magozzi & Calosi, 2015). These examples support our hypothesis, but further work using molecular approaches would be required to confirm these same mechanisms are in play in Southern Ocean diatoms in the range of temperatures relevant for observed and projected heat waves.

Analysis of a key thermal trait, T_opt_, revealed that *A.actinochilus* can alter its plastic responses to enhance growth in a warming environment, but that not all genotypes do so (Figure 5). Upward shifts in T_opt_ of up to +1.5 °C occurred in experimental populations of genotypes A4 and B7 grown at warmer acclimation temperatures and exposed to hotter heatwaves. However, we also found evidence of an upper limit to these shifts in plastic response, after which additional exposure to elevated temperatures did not increase T_opt_ further. Recent work has shown that previous environmental history can influence thermal performance, with higher acclimation temperatures resulting in upwards T_opt_ shifts for a number of phenotypic traits (Luhring & DeLong, 2017, Padfield *et al*., 2016, Seebacher *et al*., 2015). Our study, alongside others, confirms that thermal performance curves, which are often used to understand the thermal niche a taxon can occupy is plastic and can respond to rapid environmental change.

Taken together, our data show that the relative positive and negative effects of heatwaves on growth depend on thermal experience. Even though both genotype and heatwave intensity affect survival and growth, experimental populations acclimated to 2.5 °C prior to heatwave exposure generally performed better than those acclimated to 5.8 °C (Figure 3), indicating that growth temperatures above T_opt_ prior to heatwaves generally had a negative impact on growth after heatwaves. However, thermal extremes experienced during the heatwaves themselves can enhance growth rates at thermal extremes (Figure 4) and result in upwards shifts of thermal optima (Figure 5).

The thermal niches of studied Southern Ocean diatoms largely fall within the thermal annual range of the Southern Ocean (−1.5 and 8 °C) (Boyd, 2019), with thermal optima ranging between 0 and 7 °C for the majority of species (Coello-Camba & Agustí, 2017). Growth of *A. actinochilus* is consistent with this range, with thermal optimum values (3-5 °C) depending on previous thermal experience (Figure 5). The ability of populations to shift their T_opt_ towards warmer temperatures after experiencing thermal extremes may enable them to better tolerate further environmental change (Seebacher *et al*., 2015). However, the shifts in thermal responses identified in this study may be influenced by trade-offs under less permissive growth conditions, in particular under nutrient limitation. Boyd et al. (2016) investigated trait responses in the Southern Ocean diatom *Pseudonitzschia multiseries* to a number of predicted future climate scenarios. Temperature, along with iron availability, were found to be predominant drivers of phenotypic plasticity, with warmed experimental populations demonstrating enhanced growth rates and carrying capacity, but at the cost of reduced cellular quality with lower nutritional value. Their findings show that responses of SO diatoms to changing thermal environments can have important implications for trophic energy transfer and biogeochemical cycling. The enhanced growth (Figure 3) and T_opt_ shifts (Figure 5) observed in this study may well have correlated effects on ecologically important traits such as nutritional value, and future work using additional trait assays, such as determination of cellular stress and elemental analysis, could reveal them.

This study underscores the importance of intraspecific variation in heatwave responses, with genotype being both highly significant (p < 0.0001), explaining 35 % of the total variation in growth rate after heatwaves (Figure 3, supplementary table 3), and significantly affecting thermal optimum shifts after heatwaves (Figure 5, supplementary table 6). Our findings add to a growing body of literature showing that intraspecific variation in responses to environmental change is common in phytoplankton, and that multiple genotypes should be included in studies that aim to understand species-level or general responses (Godhe & Rynearson, 2017, Pancic *et al*., 2015, Schaum *et al*., 2013, Wolf *et al*., 2018, Zhang *et al*., 2014). Focus on a single genotype in this study would have produced misleading conclusions about the response of *A. actinochilus* to marine heatwaves, with either over estimation (e.g. A4) or under estimation (e.g. D8) of thermal tolerance. Our findings show that marine heatwaves can influence population dynamics and, through differential effects on lineage survival during and growth rates after heatwaves, have the potential to result in rapid adaptation in genetically-diverse diatom populations.

## CONCLUSIONS

We measured the influence of thermal experience on the survival and post-heatwave growth of a Southern Ocean diatom exposed to simulated heatwaves. We found that acclimation temperatures below T_opt_ enabled higher survival in experimental populations than did acclimation temperatures above T_opt_, which supports the hypothesis that previous thermal stress exacerbates heatwave stress. However, determination of changes in thermal response curves and T_opt_ after heatwaves that varied in both duration and temperature exposed a complex relationship between thermal experience prior to heatwaves, heatwave duration and temperature (heatwave cumulative intensity), and genetic identity. While heatwaves of severe intensity did result in drastic declines in growth rates in some genotypes, others demonstrated remarkable resistance to heatwave exposure. Furthermore, heatwaves often enhanced growth rates at the extremes of the temperature range investigated here. Both acclimation and heatwave temperatures influenced genotype-specific shifts in a key thermal trait, T_opt_, with increases of up to +1.5 °C in post-heatwave experimental populations, highlighting that both the thermal experience of experimental populations prior to heatwaves, as well as the intensity of heatwaves themselves, affect temperature response curves, which are commonly used to define the fundamental thermal niche of genotypes. Finally, heatwave survival and shifts in thermal performance caused by heatwaves differed between genotypes, indicating that there may be potential for genotype sorting and thus rapid adaptation prior to and during heatwaves in genetically-diverse diatom populations.

## Supporting information

Supplementary information - all tables and figures

## AUTHOR CONTRIBUTIONS

TR and SC obtained research funding. TS, SC and TR conceptualised the study and designed the experiments. TS performed the experiments and the data analysis, and wrote the first draft of the manuscript. SC and TR contributed to final manuscript preparation, and all authors approved the manuscript before submission.

## ACKNOWLEDGEMENTS

This work was supported by an NSF GEO-NERC grant to T.Rynearson (NSF) and S. Collins (NERC). We would like to thank Stephanie Anderson for assisting with the design of the multi-well growth plate, S. Anderson and Kerry Whittaker for cell isolations during the 2016-2017 research cruise, Celia Gelfman for culture maintenance at URI, Katy McDonald, Jana Hinners and Ignacio Melero Jiménez for assistance in data collection, Ian Bishop for providing R code for growth rate calculations and, Albert Phillimore and Nick Colegrave for advice on statistical approaches.

