## Supplementary information - all tables and figures for "Surviving heatwaves: thermal experience predicts life and death in a Southern Ocean diatom"

### Surviving Heatwaves: supplementary tables and figures

**Supplementary table 1.** Effect sizes ($\chi^2$-values), degrees of freedom and significance (p-values) of terms within the statistical model used to explain variation in the percentage of live cells remaining during heatwave exposure in the heatwave treatment. Terms in bold were statistically significant at $p<0.05$.

| Term | Effect Size ($\chi^2$-value) | df | Significance (p-value) |
|---|---|---|---|
| **Cumulative heatwave intensity** | **12.3685** | **1** | **0.0004** |
| Acclimation temperature | 3.4029 | 1 | 0.0651 |
| Genotype | 1.8158 | 2 | 0.4034 |
| Initial percentage of live cells before heatwave | 0.8001 | 1 | 0.7772 |

**Supplementary table 2.** Effect sizes ($\chi^2$-values), degrees of freedom and significance (p-values) of terms within the statistical model used to explain variation in the percentage of live cells remaining during heatwave exposure in the control treatment. Terms in bold were statistically significant at $p<0.05$.

| Term | Effect Size ($\chi^2$-value) | df | Significance (p-value) |
|---|---|---|---|
| **Acclimation temperature** | **6.4280** | **1** | **0.0112** |
| Genotype | 5.3872 | 2 | 0.0676 |
| Initial percentage of live cells before heatwave | 0.3790 | 1 | 0.5381 |
| Cumulative heatwave intensity | 0.3944 | 1 | 0.5300 |

**Supplementary table 3.** Effect sizes (F-values), degrees of freedom and significance (p-values) of terms within the statistical model used to explain variation in growth rate before and after heatwave exposure. Terms in bold were statistically significant at p<0.05.

| Term | Effect size (F-value) | df | Significance (p-value) |
|---|---|---|---|
| **Genotype** | **239.8280** | **2** | **<0.0001** |
| **Acclimation temperature (acclim_temp)** | **133.9460** | **1** | **<0.0001** |
| **Quadratic growth temperature term** | **76.0014** | **1** | **<0.0001** |
| **Linear growth temperature term (temp)** | **71.7293** | **1** | **<0.0001** |
| **Cumulative heatwave intensity (intensity)** | **67.8707** | **4** | **<0.0001** |
| **Genotype*temp** | **56.2742** | **2** | **<0.0001** |
| **Intensity*temp** | **17.7256** | **4** | **<0.0001** |
| **Genotype*acclim_temp** | **12.8787** | **2** | **<0.0001** |
| **Intensity*acclim_temp*temp** | **5.4162** | **4** | **0.0003** |
| **Genotype*temp*acclim_temp** | **5.1003** | **2** | **0.0063** |
| **Genotype*temp*intensity** | **4.3211** | **8** | **<0.0001** |
| **Genotype*temp*acclim_temp*intensity** | **2.8569** | **8** | **0.0040** |
| **Acclim_temp*intensity** | **2.7033** | **4** | **0.0297** |
| **Genotype*acclim_temp*intensity** | **2.2671** | **8** | **0.0251** |
| **Genotype*intensity** | **2.0716** | **8** | **0.0215** |
| Acclim_temp*temp | 1.5554 | 1 | 0.2128 |

**Supplementary table 4a.** Genotype A4 post-hoc pairwise comparisons between cumulative heatwave intensities (°C d) for growth rates from across the entire thermal range (-2.2 to 7.6 °C), calculated from the model presented within supplementary table 3. Note that negative estimate values indicate that growth rate is greater for the second cumulative heatwave intensity within the pair. Pairwise comparisons that were significant at p=0.05 are highlighted in bold.

| Cumulative intensity comparison | Acclimation temperature (°C) | Estimate | Standard error | df | T ratio | p-value |
|---|---|---|---|---|---|---|
| 0 - 21.87 | 2.5 | -0.03754 | 0.017642 | 629.639 | -2.12808 | 0.20941 |
| **0 - 26.61** | **2.5** | **-0.04959** | **0.016586** | **615.8426** | **-2.99006** | **0.024143** |
| 0 - 65.61 | 2.5 | -0.02779 | 0.018879 | 607.3043 | -1.472 | 0.581133 |
| 0 - 79.83 | 2.5 | 0.049872 | 0.018879 | 607.3043 | 2.541657 | 0.064287 |
| 21.87 - 26.61 | 2.5 | -0.01205 | 0.017642 | 629.639 | -0.68296 | 0.960126 |
| 21.87 - 65.61 | 2.5 | 0.009753 | 0.016856 | 623.3177 | 0.578608 | 0.978188 |
| **21.87 - 79.83** | **2.5** | **0.087416** | **0.016856** | **623.3177** | **5.186075** | **<0.0001** |
| 26.61 - 65.61 | 2.5 | 0.021802 | 0.018879 | 607.3043 | 1.154794 | 0.777039 |
| **26.61 - 79.83** | **2.5** | **0.099464** | **0.018879** | **607.3043** | **5.268453** | **<0.0001** |
| **65.61 - 79.83** | **2.5** | **0.077663** | **0.016586** | **615.8426** | **4.682549** | **<0.0001** |
| 0 - 21.87 | 5.8 | -0.03478 | 0.016586 | 615.8426 | -2.09725 | 0.222505 |
| 0 - 26.61 | 5.8 | -0.03577 | 0.017642 | 629.639 | -2.02783 | 0.253823 |
| 0 - 65.61 | 5.8 | -0.04306 | 0.018879 | 607.3043 | -2.28083 | 0.152484 |
| 0 - 79.83 | 5.8 | 0.025895 | 0.018879 | 607.3043 | 1.371625 | 0.646136 |
| 21.87 - 26.61 | 5.8 | -0.00099 | 0.017642 | 629.639 | -0.05615 | 0.999998 |
| 21.87 - 65.61 | 5.8 | -0.00828 | 0.018879 | 607.3043 | -0.43838 | 0.992324 |
| **21.87 - 79.83** | **5.8** | **0.060679** | **0.018879** | **607.3043** | **3.214076** | **0.011984** |
| 26.61 - 65.61 | 5.8 | -0.00729 | 0.016856 | 623.3177 | -0.43224 | 0.992728 |
| **26.61 - 79.83** | **5.8** | **0.06167** | **0.016856** | **623.3177** | **3.658663** | **0.002542** |
| **65.61 - 79.83** | **5.8** | **0.068956** | **0.016586** | **615.8426** | **4.15757** | **0.000353** |

**Supplementary table 4b.** Genotype B7 post-hoc pairwise comparisons between cumulative heatwave intensities (°C d) for growth rates from across the entire thermal range (-2.2 to 7.6 °C), calculated from the model presented within supplementary table 3. Note that negative estimate values indicate that growth rate is greater for the second cumulative heatwave intensity within the pair. Pairwise comparisons that were significant at p=0.05 are highlighted in bold.

| Cumulative intensity comparison | Acclimation temperature (°C) | Estimate | Standard error | df | T ratio | p-value |
|---|---|---|---|---|---|---|
| **0 - 21.87** | **2.5** | **-0.05333** | **0.016586** | **615.8426** | **-3.21558** | **0.011918** |
| **0 - 26.61** | **2.5** | **-0.05963** | **0.016586** | **615.8426** | **-3.59518** | **0.003214** |
| **0 - 65.61** | **2.5** | **-0.06439** | **0.018879** | **607.3043** | **-3.41051** | **0.006201** |
| 0 - 79.83 | 2.5 | 0.027312 | 0.018879 | 607.3043 | 1.446685 | 0.597631 |
| 21.87 - 26.61 | 2.5 | -0.0063 | 0.016586 | 615.8426 | -0.3796 | 0.995588 |
| 21.87 - 65.61 | 2.5 | -0.01106 | 0.018879 | 607.3043 | -0.58559 | 0.977199 |
| **21.87 - 79.83** | **2.5** | **0.080645** | **0.018879** | **607.3043** | **4.271603** | **0.000218** |
| 26.61 - 65.61 | 2.5 | -0.00476 | 0.018879 | 607.3043 | -0.25211 | 0.999109 |
| **26.61 - 79.83** | **2.5** | **0.08694** | **0.018879** | **607.3043** | **4.605083** | **<0.0001** |
| **65.61 - 79.83** | **2.5** | **0.0917** | **0.016586** | **615.8426** | **5.528914** | **<0.0001** |
| 0 - 21.87 | 5.8 | -0.03461 | 0.016856 | 623.3177 | -2.05303 | 0.242148 |
| 0 - 26.61 | 5.8 | -0.00471 | 0.016586 | 615.8426 | -0.28372 | 0.998582 |
| 0 - 65.61 | 5.8 | -0.01952 | 0.018879 | 607.3043 | -1.034 | 0.83954 |
| 0 - 79.83 | 5.8 | 0.040577 | 0.018879 | 607.3043 | 2.14928 | 0.200783 |
| 21.87 - 26.61 | 5.8 | 0.0299 | 0.016856 | 623.3177 | 1.773864 | 0.389819 |
| 21.87 - 65.61 | 5.8 | 0.015084 | 0.017642 | 629.639 | 0.855037 | 0.913004 |
| **21.87 - 79.83** | **5.8** | **0.075182** | **0.017642** | **629.639** | **4.261594** | **0.000226** |
| 26.61 - 65.61 | 5.8 | -0.01482 | 0.018879 | 607.3043 | -0.78475 | 0.935012 |
| 26.61 - 79.83 | 5.8 | 0.045282 | 0.018879 | 607.3043 | 2.398527 | 0.117057 |
| **65.61 - 79.83** | **5.8** | **0.060098** | **0.016586** | **615.8426** | **3.623506** | **0.002898** |

**Supplementary table 4c.** Genotype B7 post-hoc pairwise comparisons between cumulative heatwave intensities (°C d) for growth rates from across the entire thermal range (-2.2 to 7.6 °C), calculated from the model presented within supplementary table 3. Note that negative estimate values indicate that growth rate is greater for the second cumulative heatwave intensity within the pair. Pairwise comparisons that were significant at p=0.05 are highlighted in bold.

| Cumulative intensity comparison | Acclimation temperature (°C) | Estimate | Standard error | df | T ratio | p-value |
|---|---|---|---|---|---|---|
| 0 - 21.87 | 2.5 | -0.03858 | 0.016856 | 623.3177 | -2.28866 | 0.149868 |
| 0 - 26.61 | 2.5 | -0.04187 | 0.016856 | 623.3177 | -2.48376 | 0.095607 |
| 0 - 65.61 | 2.5 | -0.01807 | 0.018879 | 607.3043 | -0.95706 | 0.874121 |
| **0 - 79.83** | **2.5** | **0.105888** | **0.018879** | **607.3043** | **5.608692** | **<0.0001** |
| 21.87 - 26.61 | 2.5 | -0.00329 | 0.016586 | 615.8426 | -0.19828 | 0.999655 |
| 21.87 - 65.61 | 2.5 | 0.020509 | 0.017642 | 629.639 | 1.162513 | 0.772743 |
| **21.87 - 79.83** | **2.5** | **0.144465** | **0.017642** | **629.639** | **8.188785** | **<0.0001** |
| 26.61 - 65.61 | 2.5 | 0.023797 | 0.017642 | 629.639 | 1.348923 | 0.660601 |
| **26.61 - 79.83** | **2.5** | **0.147754** | **0.017642** | **629.639** | **8.375195** | **<0.0001** |
| **65.61 - 79.83** | **2.5** | **0.123956** | **0.016586** | **615.8426** | **7.473744** | **<0.0001** |
| **0 - 21.87** | **5.8** | **-0.07138** | **0.017642** | **629.639** | **-4.04586** | **0.00056** |
| 0 - 26.61 | 5.8 | -0.01342 | 0.016586 | 615.8426 | -0.80887 | 0.927899 |
| 0 - 65.61 | 5.8 | -0.03613 | 0.018879 | 607.3043 | -1.91389 | 0.310957 |
| **0 - 79.83** | **5.8** | **0.139624** | **0.018879** | **607.3043** | **7.395629** | **<0.0001** |
| **21.87 - 26.61** | **5.8** | **0.057961** | **0.017642** | **629.639** | **3.285414** | **0.009461** |
| 21.87 - 65.61 | 5.8 | 0.035243 | 0.016856 | 623.3177 | 2.090876 | 0.225255 |
| **21.87 - 79.83** | **5.8** | **0.211** | **0.016856** | **623.3177** | **12.51793** | **<0.0001** |
| 26.61 - 65.61 | 5.8 | -0.02272 | 0.018879 | 607.3043 | -1.20329 | 0.749505 |
| **26.61 - 79.83** | **5.8** | **0.153039** | **0.018879** | **607.3043** | **8.10623** | **<0.0001** |
| **65.61 - 79.83** | **5.8** | **0.175756** | **0.016586** | **615.8426** | **10.59696** | **<0.0001** |

**Supplementary table 5.** Effect sizes (F-values), degrees of freedom and significance (p-values) of terms within the statistical model used to explain variation in growth rate before and after heatwave exposure, without data from the lowest growth temperature (-2.24 °C). Terms in bold were statistically significant at p<0.05. The order of the terms matches the order of supplementary table 3, in which the lowest growth temperature was included within the analysed data.

| Term | Effect size (F-value) | df | Significance (p-value) |
|---|---|---|---|
| **Genotype** | **84.9725** | **2** | **<0.0001** |
| **Acclimation temperature (acclim_temp)** | **77.2271** | **1** | **<0.0001** |
| **Quadratic growth temperature term** | **99.3750** | **1** | **<0.0001** |
| **Linear growth temperature term (temp)** | **75.3006** | **1** | **<0.0001** |
| **Cumulative heatwave intensity (intensity)** | **40.5534** | **4** | **<0.0001** |
| **Genotype*temp** | **18.8273** | **2** | **<0.0001** |
| **Intensity*temp** | **12.4150** | **4** | **<0.0001** |
| **Genotype*acclim_temp** | **4.7130** | **2** | **0.0094** |
| **Intensity*acclim_temp*temp** | **4.2904** | **4** | **0.0020** |
| **Genotype*temp*acclim_temp** | **6.8828** | **2** | **0.0011** |
| Genotype*temp*intensity | 1.2177 | 8 | 0.2862 |
| **Genotype*temp*acclim_temp*intensity** | **2.0480** | **8** | **0.0393** |
| Acclim_temp*intensity | 1.7827 | 4 | 0.1309 |
| Genotype*acclim_temp*intensity | 1.6406 | 8 | 0.1106 |
| **Genotype*intensity** | **2.4793** | **8** | **0.0120** |
| Acclim_temp*temp | 0.6174 | 1 | 0.4324 |

**Supplementary table 6.** Effect sizes (F-values), degrees of freedom and significance (p-values) of terms within the statistical model used to explain variation in thermal optimum before and after heatwave exposure. Terms in bold were statistically significant at p<0.05.

| Term | Effect Size (F-value) | df | Significance (p-value) |
|---|---|---|---|
| **Cumulative heatwave intensity (intensity)** | **12.0784** | **4** | **<0.0001** |
| **Genotype** | **8.0824** | **2** | **<0.0001** |
| **Genotype*acclim_temp** | **5.995** | **2** | **0.0041** |
| **Genotype*intensity** | **2.1874** | **8** | **0.0402** |
| Acclimation temperature (acclim_temp) | 0.4319 | 1 | 0.5134 |

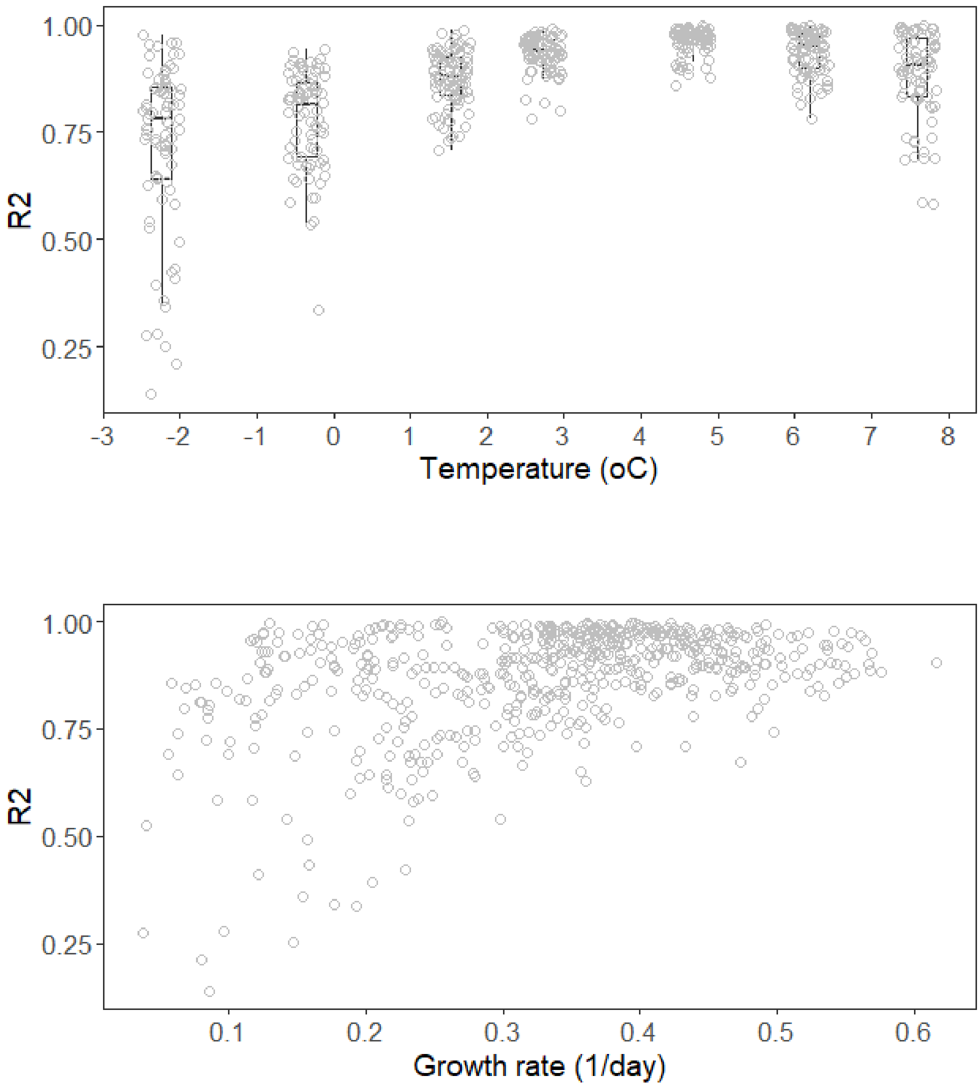


**Supplementary Figure 1.** The relationship of TPC growth rate fit $R^2$ values with TPC growth temperatures (top) and TPC growth rates (bottom).

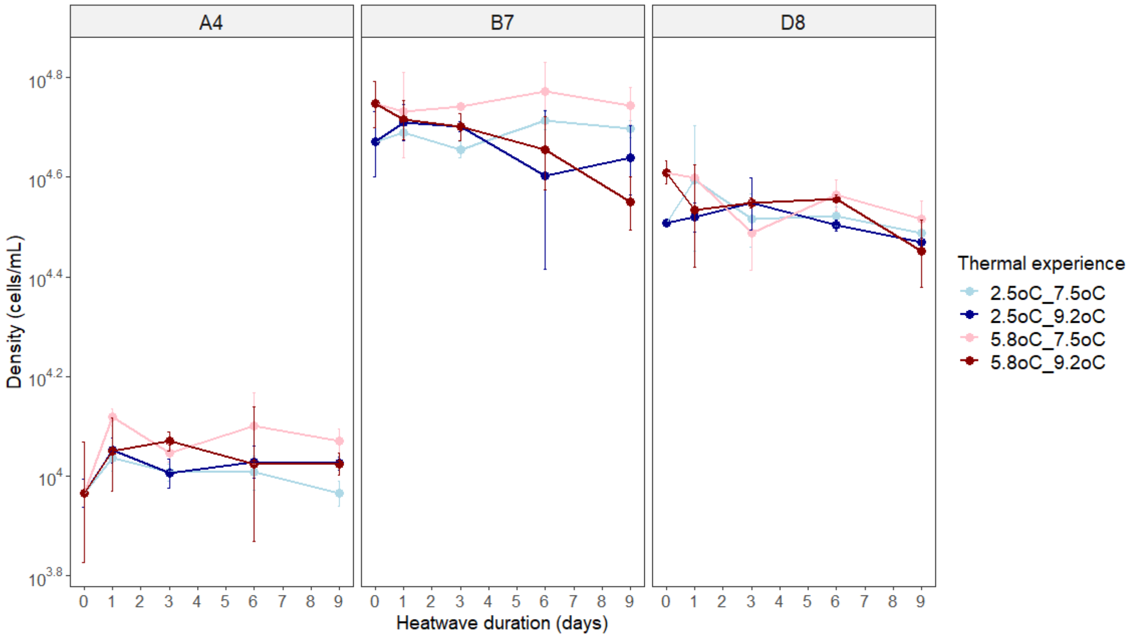


**Supplementary Figure 2.** Density (cells/mL, total live and dead cells) of experimental populations sampled after 0, 1, 3, 6 and 9 days of heatwave exposure in the heatwave treatment. Samples (1 mL volumes in 48-well microtitre plates) for each treatment were aliquoted from three biological replicates from the acclimation phase of the experiment. Data points represent average values across three biological replicates (single technical replication of cell counts), with error bars representing ± one standard deviation.

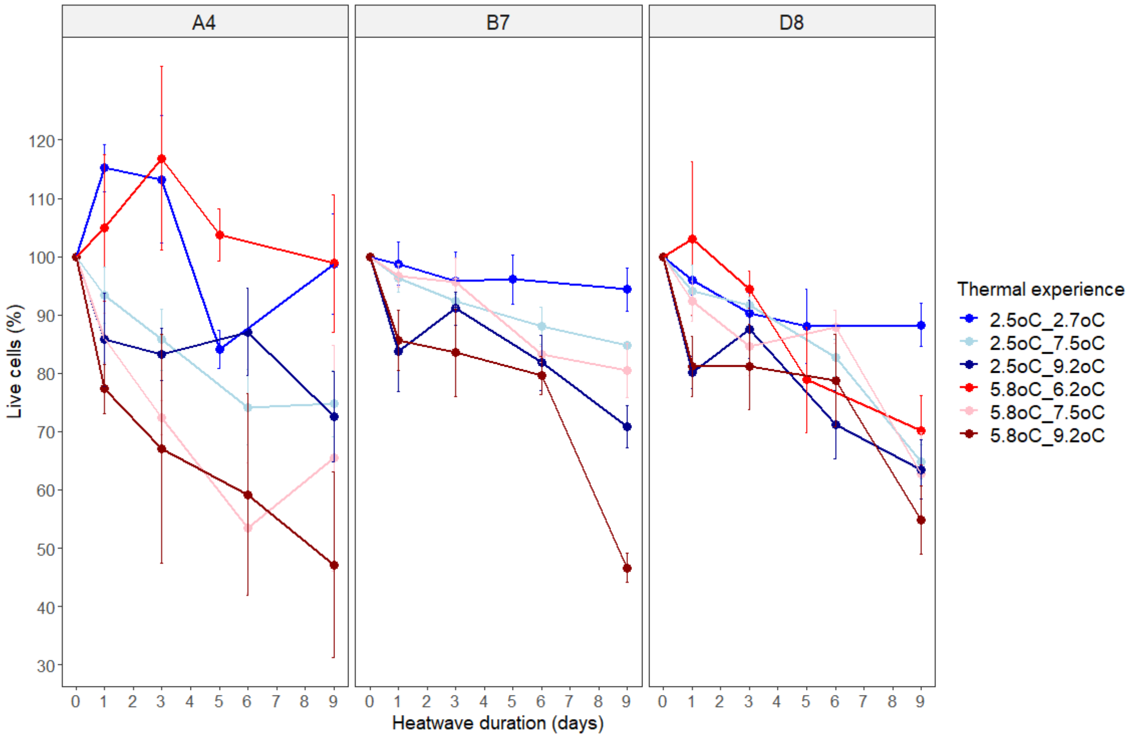


**Supplementary Figure 3.** Survival of experimental populations exposed to heatwaves in the control treatment, displayed alongside the data from the heatwave treatment. The percentage of live cells (normalised by day 0 values) were tracked across heatwave duration (0, 1, 3, 5/6 and 9 days). Thermal experience is comprised of an acclimation phase (2.5 or 5.8 °C) and a heatwave phase (2.7/6.2, 7.5 or 9.2 °C). Each panel represents the results for each of the three genotypes (A4, B7 and D8) of *Actinocyclus actinochilus*. Experimental populations of genotype B7 failed to grow at 5.8 °C in the acclimation phase of the control treatment, and as such this data is missing from this analysis. Data points represent average values across three replicates (single technical replication of cell counts), with error bars representing ± one standard deviation.

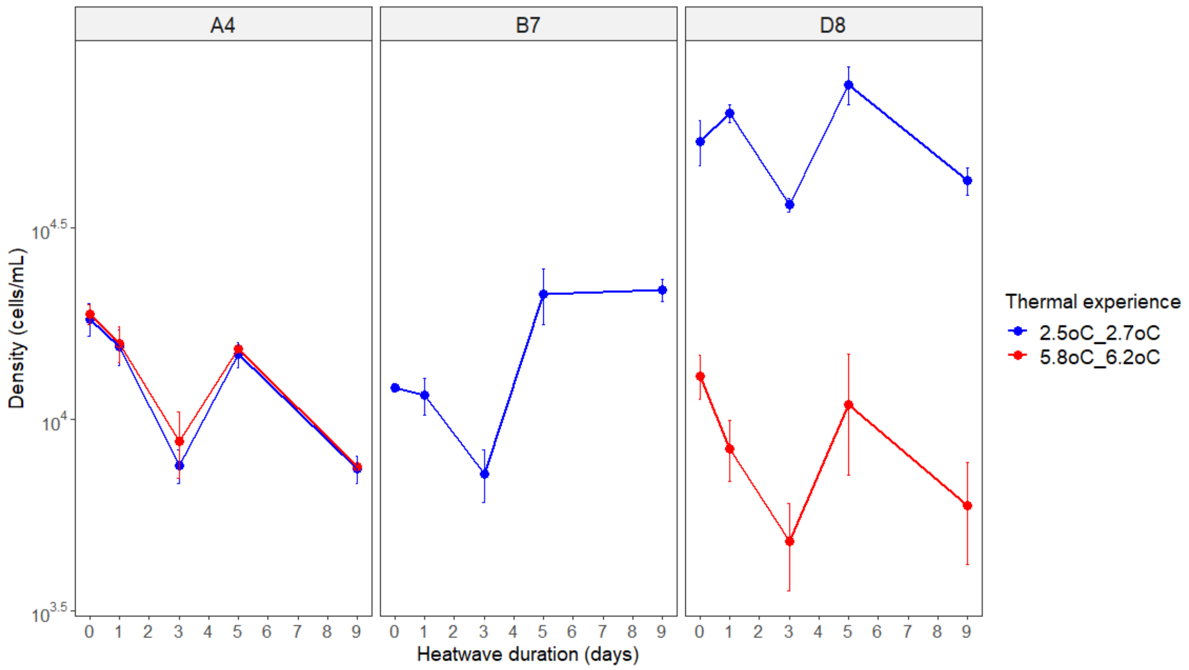


**Supplementary Figure 4.** Density (cells/mL, total live and dead cells) of experimental populations sampled after 0, 1, 3, 5 and 9 days of heatwave exposure in the control treatment. Samples (1 mL volumes in 48-well microtitre plates) for each treatment were aliquoted from three biological replicates from the acclimatory phase of the experiment. Data points represent average values across three biological replicates (single technical replication of cell counts), with error bars representing ± one standard deviation.
